# Towards development of serum substitute medium to induce osteoclast differentiation of human peripheral blood derived monocytes

**DOI:** 10.1101/2023.01.18.524526

**Authors:** Sana Ansari, Keita Ito, Sandra Hofmann

**Author notes:** Corresponding author: Sandra Hofmann.

## Abstract

Fetal bovine serum (FBS) is a widely used supplement in cell culture media despite its known drawbacks, including ethical, safety, and scientific issues. To overcome the drawbacks of using FBS in cell culture, a defined serum substitute medium needs to be developed. The development of such a medium depends on the cell type, which makes it impossible to use one universal serum substitute medium for all cells. Osteoclasts are large, multinucleated cells originated from the hematopoietic stem cell lineage that play an important role in regulating bone mass and quality. To date, no defined serum substitute medium formulations have been reported for osteoclast differentiation of monocytes derived from peripheral blood mononuclear cells (PBMCs). Here, we have attempted to develop such a serum substitute medium for the osteoclastogenesis process in a stepwise approach. Essential components were added to the medium while monocytes were cultured in 96-well plates and in Osteo-Assay well plates to analyze the formation of tartrate resistant acid phosphatase (TRAP) expressing multinucleated osteoclasts with distinct actin ring and to analyze the resorption activity of mature osteoclasts for 21 days, respectively. The serum substitute medium was aimed at supporting monocyte and later osteoclast survival, differentiation of monocytes towards multinucleated osteoclasts, and the resorption of mineralized matrix as a measure of functionality. All points were achieved after 21 days of culture in the developed serum substitute medium. This serum substitute medium could potentially replace FBS in osteoclastogenesis studies eliminating its debated use. Moreover, the well-defined serum substitute environment simplifies the study of factors released by the cells that were so far overwhelmed by the complexity of FBS.

## 1 Introduction

Fetal bovine serum (FBS) is a cell culture supplement containing hormones, growth factors, attachment factors, protease inhibitors, vitamins, and proteins that support cell survival, cell adhesion, and cell growth (Figure 1) [1]. FBS is easily accessible, inexpensive in production, and effective for most types of human and animal cells, which makes it a widely and commonly used supplement in *in vitro* cell/tissue culture experiments [2]. However, the use of FBS in the culture medium bears several drawbacks: 1) ethical aspects regarding the collection of blood from bovine fetuses, 2) biosafety aspects as FBS might contain endotoxins or viral contaminants, 3) shortage in global supply which might not meet the global demand of FBS in the future, and 4) scientific aspects as FBS is a mixture of components with qualitative, quantitative, geographical, and seasonal batch-to-batch variations. The last issue could be the reason for irreproducible and unexpected experimental outcomes within and between research groups [2], [3]. To overcome the disadvantages of using FBS in culture, defined serum substitute media with known components should be developed and replace FBS in culture [4].

**Figure 1.**
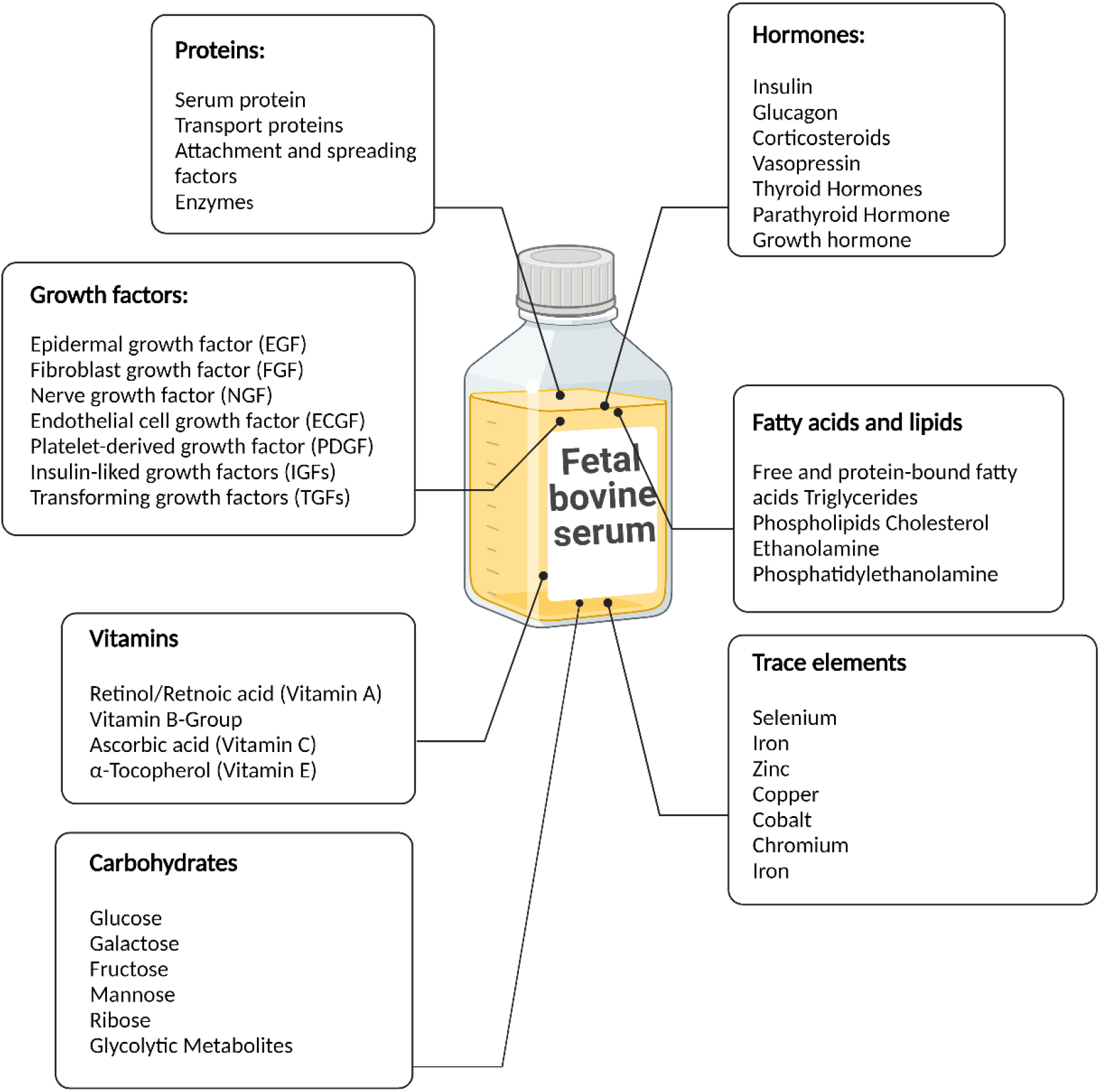
Components of FBS [5]. Created with BioRender.com.

Using serum substitute medium in *in vitro* studies holds many advantages such as: 1) avoid suffering for cows and their fetuses, 2) reduced variability in cell culture composition to obtain reliable/reproducible results, 3) support cell homeostasis in the physiological state of a specific tissue, and 4) recreate the non-physiological state of a specific tissue to study possible treatments [6]–[9]. Development of a defined serum substitute medium for cell/tissue cultures has been the focus of research for decades [4]. The identification of factors essential for cells out of many unknown and complex components of FBS is the main challenge to develop a specialized serum substitute medium which is a timely and costly process. Recently, it has been found that FBS can be replaced in the medium of some cell types, such as mesenchymal stromal cells (MSCs), with addition of certain components to maintain cell survival, growth, and differentiation potential [10], [11]. The development of these defined media depends on many factors including cell type, cell sources, applications, and species. Thus, it has been proposed that development of a universal serum substitute medium is too challenging and that the focus should be on developing a suitable one for each cell type and/or application first [4].

Osteoclasts are large, multinucleated cells originating from the hematopoietic stem cell lineage and playing a vital role in the regulation of bone mass and quality [12]. During balanced bone remodeling, osteoclast progenitor cells are recruited to the site of remodeling from the bloodstream or bone marrow and differentiate towards osteoclasts through the major influence of two cytokines: macrophage colony-stimulating factor (M-CSF) and receptor activator of nuclear factor kappa-B ligand (RANKL) which, *in vivo*, are produced by osteoblasts, osteocytes, and stromal cells [12]. These factors bind to colony-stimulating factor-1 receptor (c-fms) and receptor activator of nuclear factor kappa-B (RANK), respectively, which are the receptors on the surface of osteoclast progenitor cells. These progenitor cells fuse together to form mature multinucleated osteoclasts. Activated osteoclasts secrete hydrogen ions and proteolytic enzymes which degrade the inorganic and organic components of bone [13]–[15].

*In vitro* studies on osteoclast differentiation, activity, and their interaction with other cell types such as osteoblasts make use of cell culture medium containing FBS [16]–[19]. Studies on osteoclast differentiation in a serum substitute medium are still in their infancy probably due to the limited availability of osteoclasts, their complex differentiation process, high donor variability, and short lifespan which make studying osteoclasts challenging in general [20]–[22]. It has been reported that FBS can be replaced with specific components which could support osteoclastic differentiation in a murine monocytic cell line, RAW 264.7, but not in human monocytes derived from peripheral blood mononuclear cells (PBMCs) [23]. Thus, there is still a need to develop a serum substitute medium for osteoclast differentiation of human PBMSCs derived monocytes in a defined serum substitute medium.

The serum substitute medium was developed in a stepwise approach by adding factors to stimulate monocyte fusion and osteoclast differentiation in a basal medium containing essential components for cell survival, differentiation, and activation (Table 1). After 3 weeks of culture, the differentiation of monocytes towards osteoclasts, their activity and expression of tartrate resistant acid phosphatase (TRAP), formation of multinucleated cells with rearrangement of actin cytoskeleton, and their functionality measured as resorption potential were studied in the newly developed serum substitute medium and compared with the results of the FBS containing medium (Figure 2). Our study shows the potential of the newly developed osteoclast specific serum substitute medium for differentiation of human PBMCs derived monocytes towards mature multinucleated osteoclasts. This specialized serum substitute medium provides the opportunity to investigate the influence of different soluble factors on osteoclast formation without the complex effect of serum.

**Table 1.** Serum substitute medium components.

| **Component** | **Impact of the components on cells** | **Reference** |
| --- | --- | --- |
| Vitronectin | Enhances attachment of cells to the substrate | [24], [25] |
| Human Serum Albumin (HSA) | Acts as antioxidant to prevent oxidative damages to cells through binding to metal ions including copper or iron | [26] |
|  | Binds to various compounds including fatty acids, amino acids, hormones, vitamins, and metal ions (such as Zinc), and transport them to the cells |  |
| Antibiotic/antimycotic | Controls the growth of bacterial and fungal contamination | [27] |
| Insulin/Transferrin/Selenium | Insulin: enhances glucose and amino acid uptake by cells, intercellular transport, and synthesis of proteins | [28]–[30] |
|  | Transferrin: transports iron to cells which is a co-factor for enzymes involved in DNA synthesis and prevent extracellular oxidation |  |
|  | Selenium: protects cells from oxidative damages by reducing the production of free radical |  |
| Glutamine | An essential amino acid for the synthesis of proteins and nucleic acids | [4], [31] |
|  | Acts as a secondary energy source for metabolism |  |
| Chemically defined lipids | Serves as structural constituents of cell membrane, energy stores, and signaling molecules | [4], [32] |
| Sodium Pyruvate | A moderate excess of sodium pyruvate enhances osteoclast differentiation as this process requires significant increase in energy consumption | [33] |
| Vitamin D | Increasing the expression of osteoclast adhesion molecule $\alpha_v\beta_3$ and RANK | [34], [35] |
| Low-density lipoprotein (LDL) | Induces cell-cell fusion and controls osteoclast formation and survival | [36], [37] |

**Figure 2.**
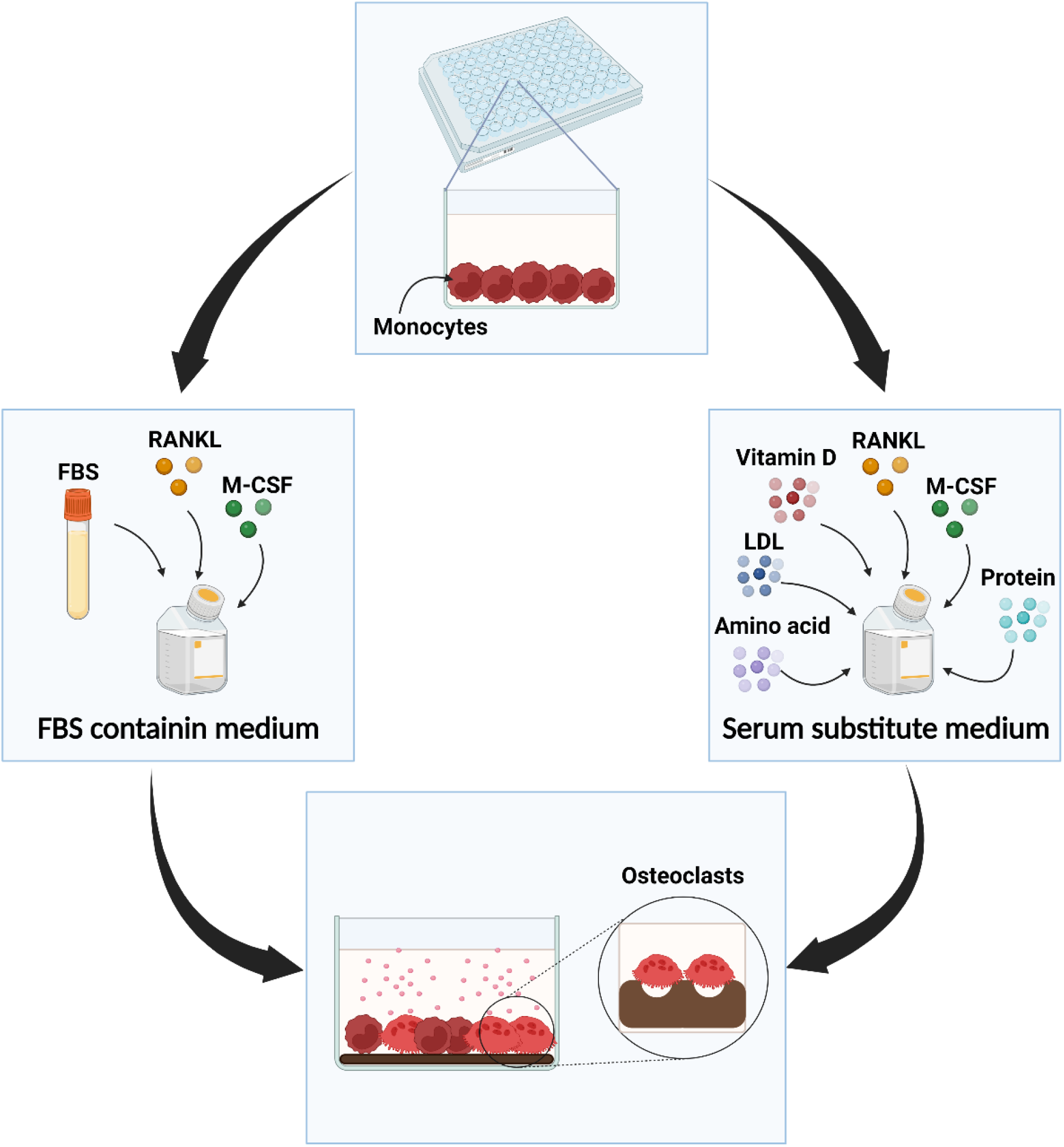
Monocytes were cultured in either FBS containing medium or serum substitute medium in which they are needed to differentiate towards TRAP-positive multinucleated osteoclasts with resorption capability. Image was created with BioRender.com.

## 2 Materials and method

### 2-1 Material

α-Minimum Essential Medium (α-MEM, 41061) and Dulbecco’s Modified Eagle Medium (DMEM, 22320) were purchased from Thermo Fisher Scientific (The Netherlands) and FBS (SFBS, Lot. No. 51113) was purchased from Bovogen (Australia). Antibiotic/antimycotic (Anti-Anti, 15240062), Insulin-Transferrin-Selenium (ITS-G, 41400045), sodium pyruvate (11360), human serum albumin (HSA, A1653), human low-density lipoprotein (LDL, LP2), and 1α,25-Dihydroxyvitamin D_3_ (vitamin D, D1530) were obtained from Sigma-Aldrich (The Netherlands). Non-essential amino acids (NEAA, 11140050), GlutaMAX (35050061), and chemically defined lipid concentrate (CD-lipid, 11905031) were from Life Technologies (The Netherlands). Basic fibroblast growth factor (b-FGF, 100-18B), recombinant human vitronectin (140-09), macrophage colony stimulating factor (M-CSF, 300-25), and receptor activator nuclear factor kappa-B ligand (RANKL, 310-01) were purchased from Peprotech (UK). Recombinant human bone morphogenetic protein-2 (rhBMP-2, 7510200) was purchased from Medtronic Sofamor Danek (USA). Unless noted otherwise, all other substances were of analytical or pharmaceutical grade and obtained from Sigma-Aldrich (The Netherlands).

### 2.2 Serum substitute medium development

In the first step of development of serum substitute medium for osteoclast cultures, we decided to use the medium that we previously developed for osteoblast cultures [38]. We aimed at investigating whether the same medium supplemented with osteoclast differentiation supplements (M-CSF and RANKL) would be sufficient for initiating osteoclastogenesis of human PBMC derived monocytes. In the next step, as the osteoblast-specific serum substitute medium supplemented with M-CSF and RANKL did not seem to be sufficient to initiate osteoclastogenesis, we aimed at developing a novel serum substitute medium specific for osteoclast cultures. To induce the formation of multinucleated mature osteoclasts, several components such as vitamin D and LDL were added to the primary serum substitute medium. Figure 3 illustrates the stepwise approach of components added.

**Figure 3.**
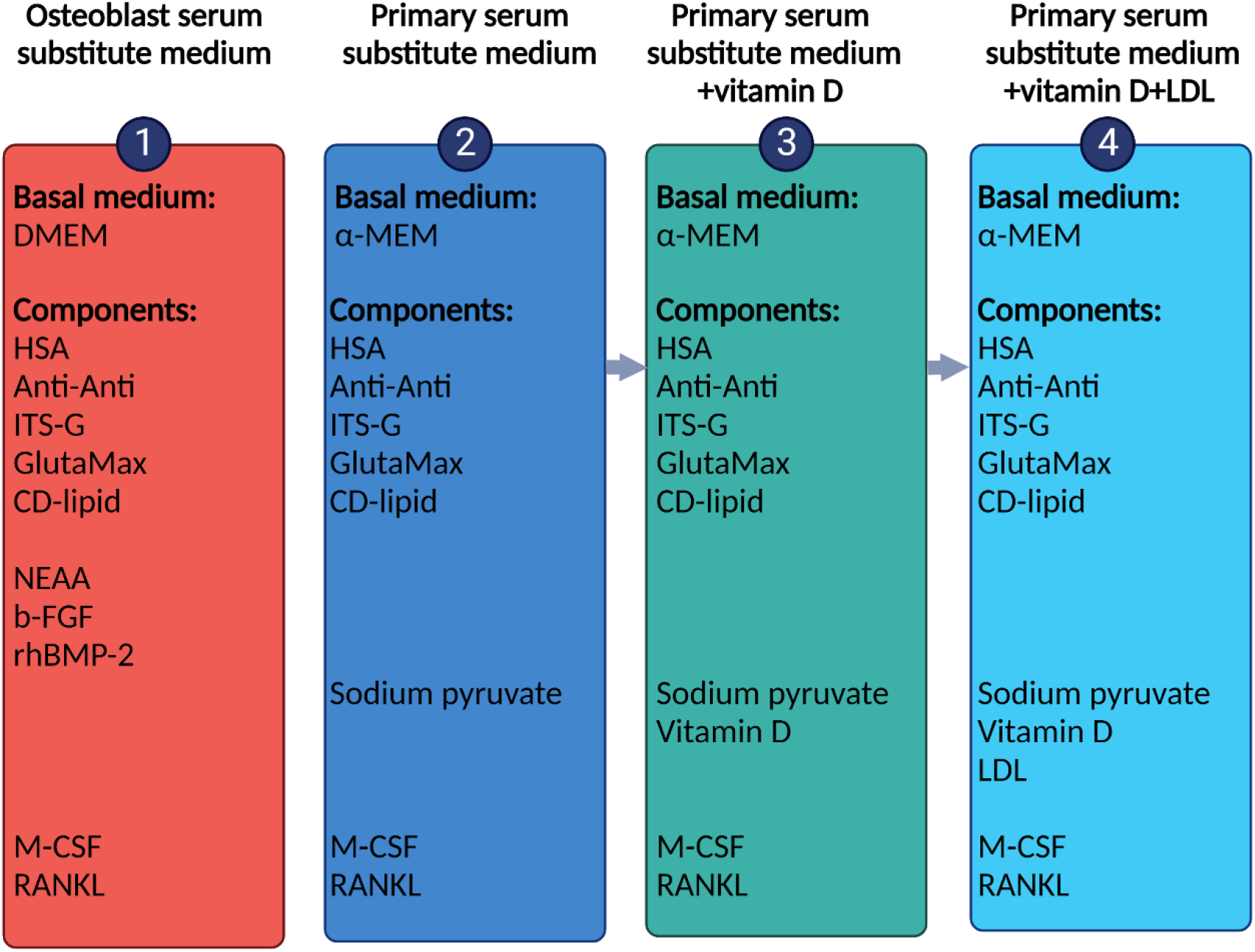
The steps of development serum substitute medium for osteoclast cultures. Image was created with BioRender.com.

#### 2.2.1 Step 1-osteoblast serum substitute medium

Initially serum substitute medium consisted of DMEM, 1% w/v HSA, 1% v/v Anti-Anti, 1% v/v ITS-G, 1% v/v GlutaMax, 1% v/v NEAA, 0.1% v/v CD-lipid, 10 ng/ml b-FGF, and 100 ng/ml rhBMP-2 and supplemented with osteogenic differentiation factors: 50 μg/mL ascorbic-acid-2-phosphate (Sigma-Aldrich, A8960), 100 nM dexamethasone (Sigma-Aldrich, D4902), 10 mM β-glycerophosphate (Sigma-Aldrich, G9422), as previously described for osteoblasts [38]. To this, osteoclast differentiation factors were added: 50 ng/mL M-CSF for the first 48 hours, and 50 ng/mL M-CSF and 50 ng/mL RANKL for the rest of the experiment [39].

#### 2.2.2 Step 2-primary serum substitute medium

To develop a serum substitute medium specifically for osteoclast cultures, we selected α-Minimum Essential Medium (α-MEM, Cat. No. 41061) as it has been shown before to best support osteoclast formation from monocytes derived from PBMCs when supplemented with 10% FBS [39], [40]. α-MEM contains a wide variety of amino acids, vitamins, inorganic salts, glucose, and sodium pyruvate that each have specific functions essential for cell function (Table 2).

**Table 2.** The components of α-MEM and their function on cells.

| Components | Impact of the components on cells | references |
| --- | --- | --- |
| Amino acids | Basic building blocks of proteins and essential for cell proliferation and viability | [41] |
| Vitamins | Essential for cell growth and proliferation, act as co-factor for enzymes | [4], [42] |
| Inorganic salts | Maintain the osmotic balance and regulate membrane potential which is necessary to transport ions through cell membrane | [43], [44] |
| Glucose | Main source of energy for cell metabolism | [27] |
| Sodium Pyruvate | By-product of glucose consumption in the glycolytic pathway and functions as a source of energy | [45] |

To induce osteoclast differentiation, the primary serum substitute medium was developed by adding 1% w/v HSA, 1% v/v Anti-Anti, 1% ITG-S, 1% v/v GlutaMAX, 0.1% v/v CD-lipid, 1% v/v sodium pyruvate to α-MEM. Again, this medium was supplemented with 50 ng/ml M-CSF for the first 48 hours, and 50 ng/mL M-CSF and 50 ng/mL RANKL for the rest of the experiment.

#### 2.2.3 Step 3-primary serum substitute medium + vitamin D

To enhance the osteoclast differentiation of monocytes, the primary serum substitute medium (2-2-2) was additionally supplemented with 20 nM vitamin D. It has been shown that vitamin D can support the formation of multinucleated osteoclasts [34], [46]. This medium was also supplemented with 50 ng/ml M-CSF for the first 48 hours, and 50 ng/mL M-CSF and 50 ng/mL RANKL for the rest of the experiment.

#### 2.2.4 Step 4- primary serum substitute medium + vitamin D + LDL

Since it seemed that cell-cell fusion was still insufficient with medium 2-2-3, 20 µg/ml LDL was additionally added. LDL has been shown to be essential for cell-cell fusion [36], [37]. Again, this medium was supplemented with 50 ng/ml M-CSF for the first 48 hours, and 50 ng/mL M-CSF and 50 ng/mL RANKL for the rest of the experiment.

#### 2.2.5 Control medium

As controls, either DMEM (for step 1) or α-MEM (for steps 2-4) were supplemented with 10% FBS, 1% Anti-Anti, and osteoclast differentiation factors: 50 ng/mL M-CSF for the first 48 hours, and 50 ng/mL M-CSF and 50 ng/mL RANKL for the rest of the experiment.

### 2.3 Monocyte isolation, seeding, and cultivation

#### Monocyte isolation

Human peripheral blood buffy coats from 2 healthy donors were collected under institutional guidelines with informed consent per declaration of Helsinki (Sanquin, Eindhoven, The Netherlands). 50 ml of buffy coats were diluted with 0.6% (w/v) sodium citrate in PBS (citrate-PBS) up to a final volume of 200 mL and layered per 25 mL on top of 10 mL Lymphoprep™ (Stem Cell Technologies, Koln, Germany). The samples were centrifuged at 800g with the lowest break for 20 minutes. PBMCs were collected, resuspended in citrate-PBS, washed 4 times in citrate-PBS supplemented with 0.01% bovine serum albumin (BSA, Roche, 10.7350.86001). PBMCs were frozen at 50*10^6 cells/ml in freezing medium containing α-MEM, 20% human platelet lysate (hPL, PE20612, PL BioScience, Aachen, Germany), and 10% dimethyl sulfoxide (DMSO, VWR, 1.02952.1000. Radnor, PA, USA) and stored in liquid nitrogen until further use. To isolate monocytes, PBMCs were thawed in α-MEM containing 10% hPL and 1% Anti-Anti, centrifuged at 300g for 10 minutes, and resuspended in isolation buffer (0.5% w/v BSA in 2 mM EDTA-PBS). Monocytes were isolated with magnetic activated cell separation (MACS) using the Pan Monocyte Isolation Kit (130-096-537, Miltenyi Biotec, Leiden, The Netherlands) and LS columns (Miltenyi Biotec, 130-042-401) according to the manufacturer’s instructions.

#### Monocyte seeding and cultivation

The wells of a 96-well plate were coated with 5 µg/mL vitronectin diluted in PBS. Briefly, 100 µL of vitronectin solution was added to each well and incubated in an incubator (37°C, 5% CO_2_) for 2 hours and then at 4°C overnight. The following day, the vitronectin solution was aspirated, the wells were rinsed 3 times with PBS, and the well-plate was pre-warmed in an incubator before seeding the cells. Monocytes were seeded in the well-plate at a density of 9*10^4^ cells per well (n=5-6). Monocytes were cultured in either **FBS containing medium** or the respective **Serum substitute medium** (section 2-2) containing 50 ng/ml M-CSF for the first 48 hours. After 48 hours, the medium was replaced with **FBS containing medium** or the respective **Serum substitute medium** containing 50 ng/ml M-CSF and 50 ng/ml RANKL to induce osteoclast differentiation. The medium was replaced 3 times per week for 3 weeks.

For Step 4, to study the resorption activity of differentiated osteoclast in FBS containing medium and serum substitute medium, monocytes of each donor were seeded in a vitronectin coated 96 Osteo-Assay 96 well-plate (CLS3988, Corning, Amsterdam, The Netherlands) at a density of 9*10^4^ cells per well (n=5-6). Monocytes were cultured in either **FBS containing medium** or **Serum substitute medium** containing 50 ng/ml M-CSF for the first 48 hours. After 48 hours, the medium was replaced with **FBS containing medium** or **Serum substitute medium** containing 50 ng/ml M-CSF and 50 ng/ml RANKL to induce osteoclast differentiation. The medium was replaced 3 times per week for 3 weeks.

### 2.4 Immunohistochemistry

To visualize cell nuclei, the actin cytoskeleton, and the expression of TRAP, immunohistochemistry was performed. Cell-seeded wells were rinsed with PBS and fixed with 10% neutral buffered formalin for 30 minutes at 4°C. Then, the wells were rinsed 3 times with PBS and covered with 100 µL 0.5% (w/v) Triton x-100 (Merck, 1.08603.1000) in PBS for 5 minutes to permeabilize cells. Then, the wells were rinsed with PBS and incubated with 5% (v/v) normal goat serum and 1% (w/v) BSA in PBS for 1 hour at room temperature to block non-specific antibody binding. Wells were then incubated overnight at 4°C with primary antibody solution containing 5% (v/v) normal goat serum, 1% (w/v) BSA in PBS, and anti-TRAP antibody (Abcam, ab185716, 1:200). The following day, the wells were washed with PBS 3 times and incubated for 1 hour with secondary antibody solution containing 5% (v/v) normal goat serum, 1% (w/v) BSA in PBS, and anti-mouse (Molecular probes, A21240, 1:200). This was followed by 3 times rinsing of the wells with PBS and incubation of cells with 0.1 µg/ml 4′,6-diamidino-2-phenylindole (DAPI (Sigma-Aldrich, D9542) and 50 pmol Atto 488-conjugated Phalloidin (Sigma-Aldrich, 49409, diluted in PBS) for 30 minutes at room temperature to stain nuclei and actin cytoskeleton, respectively. Wells were rinsed with PBS 3 times and then covered with PBS. The expression of proteins was visualized with a Leica TCS SP5X microscope and images were processed with ImageJ (version 1.53f51). Figures were chosen to be representative images per group for all the samples assessed.

### 2.5 Measurement of number of nuclei per osteoclasts and size of osteoclasts

TRAP-positive cells with 3 or more nuclei were considered to be osteoclasts. To quantify the number of nuclei per osteoclasts and size of osteoclasts, 6-10 immunohistochemistry images of cells grown in step 4 and control medium were analyzed using ImageJ (version 1.53f51). The number of nuclei per TRAP-positive cells were manually counted. The longest diameter of osteoclasts was measured using the length measurement function of ImageJ after calibrating it using the scale bar of each image.

### 2.6 Tartrate resistant acid phosphatase (TRAP) assay

The release of TRAP to the culture medium as a measure for osteoclast differentiation was measured in the cell supernatant. 10 µL supernatant or p-nitrophenol standard was incubated in 90 µL p-nitrophenyl phosphate buffer (1 mg/ml p-nitrophenyl phosphate disodium hexahydrate (Sigma-Aldrich, 71768), 0.1 M sodium acetate, 0.1% Triton x-100, and 30 µL/mL tartrate solution (Sigma-Aldrich, 3873) in PBS) in a 96-well plate for 90 minutes at 37 °C. The reaction was stopped by adding 100 µL 0.3 M NaOH. Absorbance was measured using a plate reader at 405 nm and TRAP activity was calculated by comparison to standards of known p-nitrophenol concentration.

### 2.7 Resorption assay

The resorption activity of osteoclasts was measured after 21 days of culture in FBS containing medium and complete serum substitute medium. First, cells seeded on Osteo-Assay wells were removed by incubating in 5% bleach in ultra-pure water (UPW) for 5 minutes, following rinsing the wells twice with UPW. To visualize the non-resorbed surface, the remaining calcium phosphate within the wells were stained with Von Kossa as described previously [40], [47]. Briefly, wells were incubated in 5% (w/v) silver nitrate (Sigma-Aldrich, 209139) in UPW for 30 minutes in dark, following rinsing with UPW, and incubation in 5% (w/v) sodium carbonate (Sigma-Aldrich, S7795) in 10% neutral buffered formalin for 4 minutes. The solution was completely aspirated, and the plates were dried at 50°C for 1 hour. The wells were captured with bright field microscope (Zeiss Axio Observer Z1) using tile scanning function. The tile scans were stitched with Zen Blue software (version 3.1, Zeiss). The segmentation and resorption quantification were done as previously described [40]. Briefly, first image contrast was increased using ImageJ. Then, a clipping mask was created in Illustrator (Adobe Inc., San Jose, Ca, USA) in order to remove the edges of the wells. Segmentation was done using MATLAB (version 2019b, The MathWorks Inc., Natrick, MA, USA), using Otsu’s method for binarization with global thresholding. The total number of pixels within the well and the number of resorbed pixels were determined to quantify the percentage of resorbed area.

### 2.8 Statistics

GraphPad Prism 9.0.2 (GraphPad Software, La Jolla, CA, USA) was used to perform statistical analysis and to prepare the graphs. Data used for statistical analysis was tested for normality using the Shapiro-Wilk normality test. The TRAP assay data (Figure 4C, 5D, and 6E) were normally distributed. These data were compared using Repeated Measures Two-Way Analysis of Variances (ANOVA) followed by Tukey’s post hoc tests with adjusted p-value for multiple comparisons. Geisser-Greenhouse correction was used to account for unequal variances. The data is presented as mean and standard deviation. Number of nuclei per osteoclasts (Figure 6C), size of osteoclasts (Figure 6D), and resorption data (Figure 7C) were not normally distributed and were tested with Mann-Whitney test and are presented as median and interquartile range. Differences were considered statistically significant at a level of p-value<0.05.

**Figure 4.**
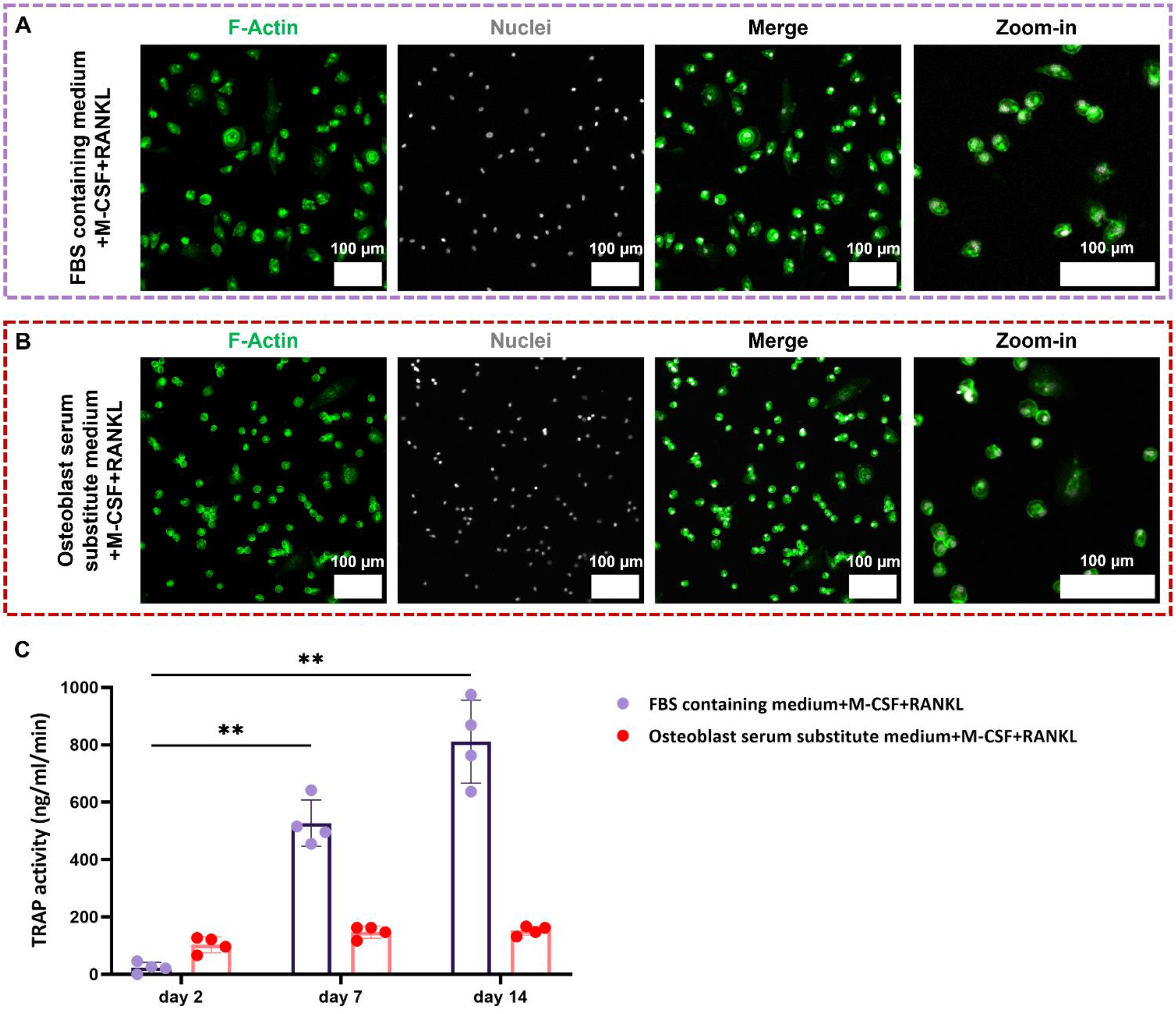
In step 1 of development serum substitute medium for osteoclast cultures, the previously developed serum substitute medium for osteoblast cultures supplemented with M-CSF and RANKL was used. Monocytes did not differentiate towards multinucleated osteoclasts in FBS containing medium (A) and osteoblast serum substitute medium (B) after 14 days of culture. TRAP release into the medium increased significantly over time in FBS containing medium (p<0.01). Only a slight increase in TRAP release was detected in osteoblast specific serum substitute medium (C).

## 3. Results

### 3.1 Serum substitute medium for osteoblast cultures containing M-CSF and RANKL did not support osteoclast formation

In the first step to develop a serum substitute medium for osteoclast cultures, we used a previously developed serum substitute medium for osteoblast cultures [38] supplemented with osteoclast differentiation factors: M-CSF and RANKL. This was done to investigate whether only adding the osteoclast differentiation factors to an already functional serum substitute medium would be sufficient for osteoclast cultures. The results indicated that monocytes did not differentiate towards multinucleated osteoclasts in both FBS containing medium (Figure 4A) and osteoblast serum substitute medium (Figure 4B). In both medium types, cells seemed to remain as mononucleated monocytes despite the presence of M-CSF and RANKL. Interestingly, TRAP release into cell culture supernatant increased significantly over time in FBS containing medium (Figure 4C), while in serum substitute medium, only a slight increase could be detected.

### 3.2 Development of serum substitute medium for osteoclastogenesis

Monocytes were cultures in the “ primary serum substitute medium” (Figure 2-Step 2) supplemented with 50 ng/ml M-CSF for the first 48 hours and 50 ng/ml M-CSF and 50 ng/ml RANKL for the next 21 days. As a control, monocytes were also cultured in FBS containing medium supplemented with the same amount of M-CSF and RANKL. Multinucleated osteoclasts were formed in FBS containing medium (Figure 5A), while in the primary serum substitute medium monocytes remained mononucleated and did not differentiate into multinucleated osteoclasts (Figure 5B). TRAP release into the cell culture supernatant increased significantly over time in the FBS containing medium (Figure 5D). The primary serum substitute medium did not induce any TRAP activity by cells (Figure 5D).

**Figure 5.**
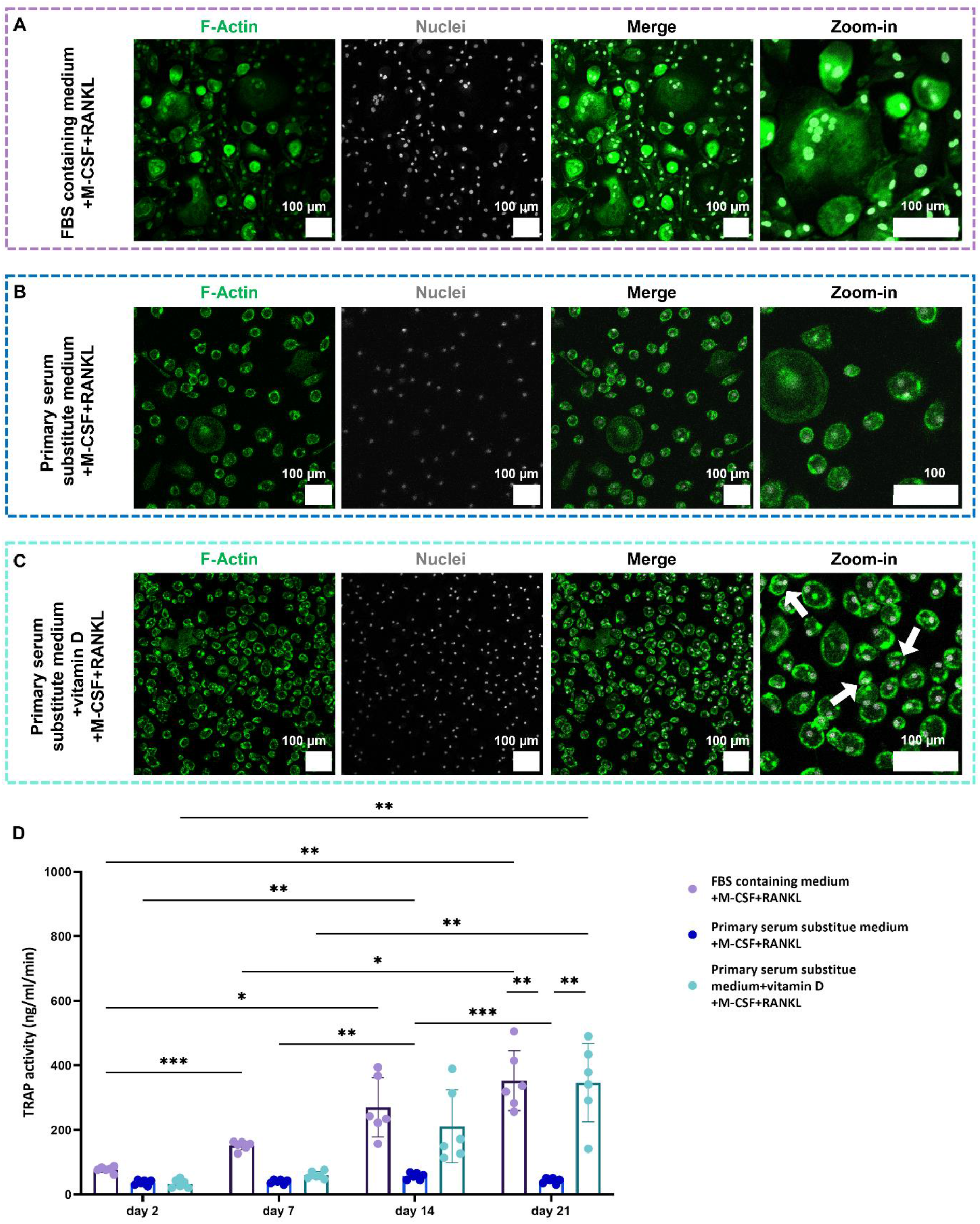
Vitamin D in serum substitute medium enhanced the formation of osteoclasts in culture. Monocytes differentiated into multinucleated osteoclasts in FBS containing medium (A). Primary serum substitute medium did not induce osteoclast formation (B). Addition of vitamin D induce formation of few double-nucleated osteoclasts (C). TRAP activity of cells cultured in FBS containing medium increased significantly over time. The primary serum substitute medium did not affect the TRAP release by cells. Addition of vitamin D to primary serum substitute medium significantly enhanced TRAP activity of cells (D). * p<0.05, ** p<0.01, *** p<0.001.

In the next step of developing a serum substitute, it was chosen to additionally supplement the primary serum substitute medium with vitamin D to support the formation of multinucleated osteoclasts [34], [46]. The addition of 20 nM vitamin D to the primary serum substitute medium supported some fusion of cells into double-nucleated cells (Figure 4C). But more often, mononucleated monocytes remained in each other’s vicinity and only in a few spots, cells did fuse together (Figure 5C, arrows). However, the addition of vitamin D was able to increase TRAP activity to an equal level as the FBS containing control group (Figure 5D). After 21 days of culture, no significant differences were detected in TRAP activity of cells cultured in FBS containing medium and primary serum substitute medium containing vitamin D.

### 3.3 Promote cell-cell fusion in serum substitute medium

Osteoclasts are characterized as large (10-300 µm in diameter), multinucleated cells with at least 3 nuclei per cell and a typical actin ring [54], [55]. In the next step, we tried to further promote the cell-cell fusion. This was done by adding low-density lipoprotein (LDL) to the primary serum substitute medium containing vitamin D. The addition of LDL led to the formation of large, multinucleated cells with a clear actin ring (Figure 6B), similar to FBS containing medium (Figure 6A). The formed multinucleated cells contained more nuclei compared to double-nucleated cells formed in the absence of LDL (Figure 6B-arrows *vs*. Figure 5C-arrows). This serum substitute medium induced the formation of larger osteoclasts with less nuclei compared to FBS containing medium in which smaller osteoclasts were formed with more nuclei per osteoclast (Figure 6C and 6D). In both FBS containing medium and serum substitute medium, round mononucleated cells were also present which suggested that not all cells differentiated towards osteoclasts. TRAP was expressed both in monocytes and osteoclasts cultured in both FBS containing medium and serum substitute medium.

**Figure 6.**
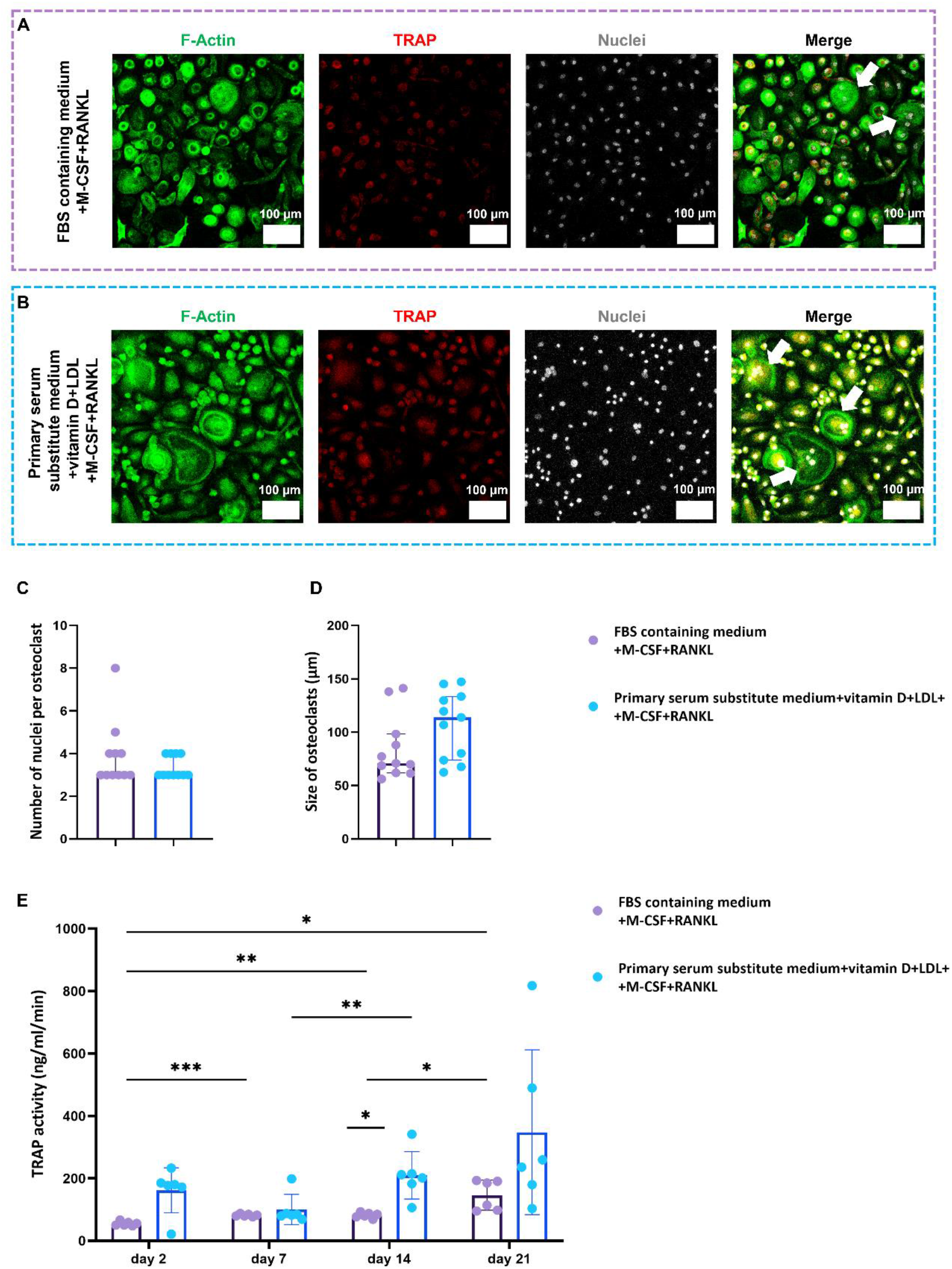
The serum substitute medium induced osteoclast differentiation. Monocytes differentiated into multinucleated osteoclasts in FBS containing medium (A). Primary substitute medium containing vitamin D and LDL supported the formation of multinucleated osteoclasts after 21 days of culture (B). *The serum substitute medium induced the formation of larger osteoclasts with less nuclei compared to FBS containing medium in which smaller osteoclasts were formed with more nuclei per osteoclast (C and D)*. TRAP activity of cells cultured in serum substitute medium increased over time in both groups (E). * p<0.05, ** p<0.01, *** p<0.001.

As expected, in FBS containing medium, TRAP release into the medium increased significantly over time. Similarly, TRAP activity of cells cultured in the serum substitute medium containing vitamin D and LDL increased over time. The variation in TRAP activity was larger compared to the FBS containing medium (Figure 5C). The release of TRAP to the serum substitute medium was in general slightly higher than in FBS containing medium, however only on day 14, the TRAP activity of cells in serum substitute medium was significantly higher than in FBS containing medium. Higher TRAP release in serum substitute medium after 14 days compared to FBS containing medium could suggest that the serum substitute medium stimulated cells to express and release TRAP to the medium faster than in FBS containing medium.

### 3.4 Serum substitute medium containing vitamin D and LDL supported osteoclast activity

The functionality of osteoclasts *in vitro* in FBS containing medium and serum substitute medium containing vitamin D and LDL were analyzed by the assessing their capacity to resorb calcium phosphate surfaces (Osteo-Assay plates). Osteoclasts formed in both FBS containing medium (Figure 7A) and serum substitute medium (Figure 7B) were able to resorb parts of the Osteo-Assay plates. The resorbed and un-resorbed area are shown in white and black, respectively. Quantification of the resorbed area of the Osteo-Assay plates by differentiated osteoclasts in FBS containing medium revealed a variation from 0.04% to 11% (Figure 7C). This variation could be attributed to a donor-dependent response of osteoclasts to FBS, which has been described earlier [40]. Osteoclasts formed in serum substitute medium were not able to resorb the whole wells (Figure 7B) and the quantification of the resorbed area of Osteo-Assay plates by differentiated osteoclasts in complete serum substitute medium showed to be 2.86 +/-1.28% (Figure 7C).

**Figure 7.**
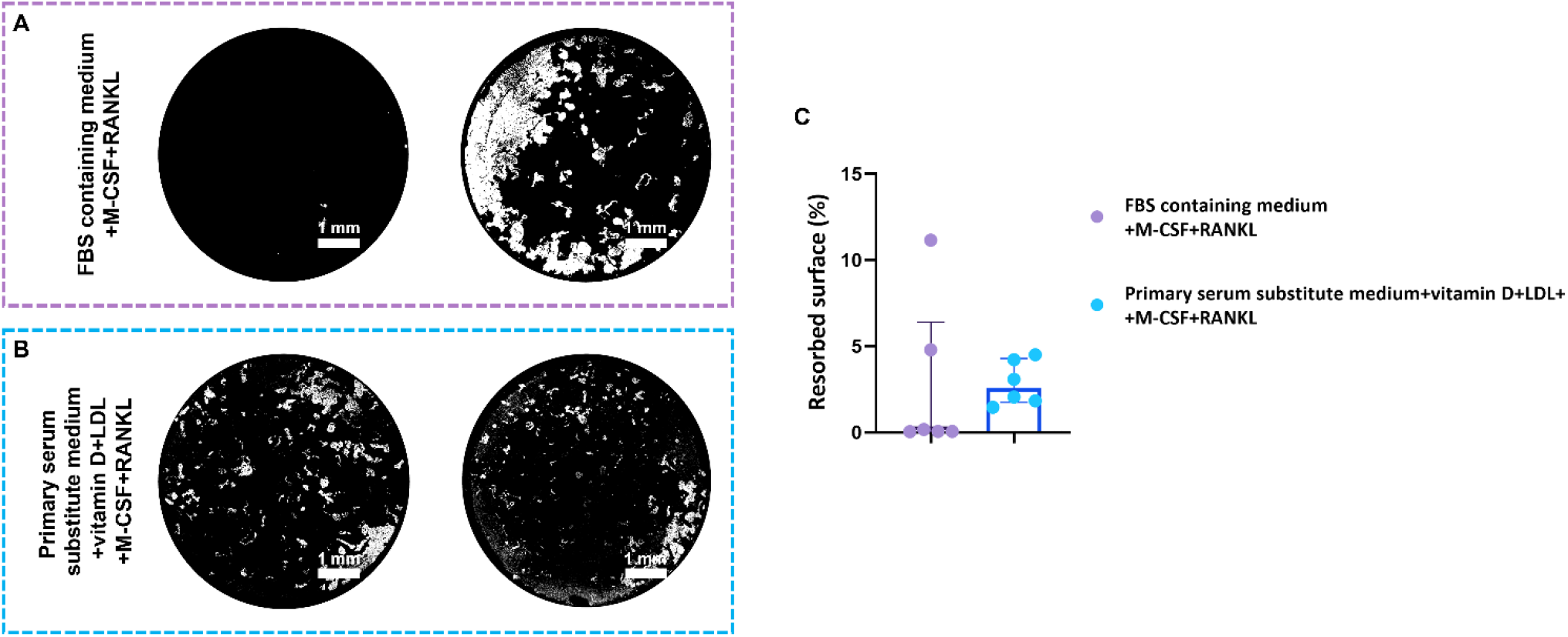
Osteoclasts formed in serum substitute medium containing vitamin D and LDL resorbed the calcium phosphate surface. The Osteo-Assay surface was resorbed in both FBS containing medium (A) and serum substitute medium (B). Quantification of resorbed area showed differences in resorption potential of differentiated osteoclasts in FBS containing medium and serum substitute medium (C).

## Discussion

FBS is a commonly used growth supplement in most cell/tissue culture media despite its known disadvantages in scientific, biosafety, and ethical aspects [2]. To overcome the disadvantages of using FBS in cell culture media, defined media replacing FBS need to be developed. Due to the differences in cell’s requirements in growth and differentiation, the defined media might need to be optimized for each specific cell type. In the present study, we developed a serum substitute medium in a step-by-step approach for osteoclastogenesis of human PBMCs derived monocytes. By doing so, we could also elucidate the effect that the added supplements have during osteoclastogenesis. The suitability of our newly developed serum substitute medium was determined based on its capability to support the differentiation of monocytes towards resorbing TRAP-positive multinucleated osteoclasts with a clear actin ring.

Osteoclasts are large multinucleated cells formed by fusion of mononuclear monocytes capable of resorbing bone. Each osteoclast contains 3 or more nuclei and the cell size varies in diameter between 10 to 300 µm [54]. Despite many attempts to develop a serum substitute medium for different cell types [3]– [5], a defined serum substitute medium for osteoclast differentiation from human monocytes has not been reported yet. This might be attributed to the challenges of studying osteoclasts in general due to their limited availability, complex differentiation process, donor variation, and short lifespan [20]–[22]. In an early study, there have been attempts to define a serum substitute medium to differentiate human monocytes and the murine monocytic cell line, RAW 264.7 towards osteoclasts [23]. This defined serum substitute medium contained albumin, transferrin, insulin, β-mercaptoethanol, LDL, ascorbic acid, epidermal growth factor (EGF), platelet derived growth factor-BB (PDGF-BB), dexamethasone, and glutamine and has been shown to support the formation of TRAP releasing multinucleated osteoclasts from RAW 264.7. However, the same medium did not support osteoclast formation of human PBMCs derived monocytes [23]. Thus, to be able to study human osteoclast differentiation in a more defined biochemical environment, there is still a need to develop a serum substitute medium for osteoclastogeneis of human monocytes.

We used a stepwise approach for serum substitute development. In the first step, a previously developed serum substitute medium for osteoblast cultures which was able to support osteogenic differentiation of MSCs and ECM deposition after 21 days of culture has been used [38]. This step was done to investigate whether the same medium supplemented with osteoclast differentiation factors (M-CSF and RANKL), which are expressed by mature osteoblasts [56], would be sufficient for osteoclast differentiation of monocytes. The potential of using the same serum substitute medium was in view of using it for studies mimicking the bone remodeling process [39]. In such bone remodeling models, bone forming osteoblasts and bone resorbing osteoclasts act closely together in a co-culture. One of the challenges in creating such models has been the choice of medium composition [39], [57]. A suitable medium for co-culture experiments needs to support the growth and function of both cell types. Thus, it would have been ideal if the same serum substitute medium developed for osteoblast cultures could also support osteoclast formation. However, here we have shown that culturing monocytes in osteoblast serum substitute medium supplemented with osteoclast differentiation factors was not able to induce osteoclast differentiation. The inability of FBS containing medium to support differentiation of monocytes towards osteoclasts could be due to the use of DMEM as basal medium. Osteoblast serum substitute medium did not induce osteoclast differentiation probably due to not only the use of DMEM as basal medium but also the presence of factors that inhibit the osteoclast formation such as dexamethasone. DMEM contains phenol red (pH indicator). Previous studies have shown that phenol red can mimic the action of estrogen and inhibit osteoclast formation and activity [48], [49], [50], [51]. Moreover, the presence of dexamethasone in high concentrations (100 nM) in osteoblast serum substitute medium has been shown to inhibit the formation of TRAP-positive multinucleated osteoclasts [52], [53]. These results indicated that serum substitute medium containing factors that support osteoblast cultures could interferes with monocyte differentiation into osteoclasts even when potent osteoclast differentiation factors are present. This observation suggested that for monocyte differentiation into osteoclasts, different formulation of serum substitute medium would be needed. In future studies, a serum substitute medium should be developed that supports both cell types in a co-culture. It might be that interaction between the cells could limit the need to add exogenous factors. For instance, the expression of M-CSF, RANKL, and osteoprotegerin (OPG) by mature osteoblasts could support osteoclast formation in a co-culture [40], [58].

In the next step, we focused on developing a specialized serum substitute medium for osteoclast differentiation from human PBMC derived monocytes. We selected α-MEM (without phenol red) as a basal medium as it has been shown before to support osteoclast formation when supplemented with 10% FBS [39], [40]. Phenol red is a pH indicator in cell culture media, but studies have shown that it mimics the action of estrogen and its effect should be taken into account when studying estrogen-responsive cells such as osteoclasts [48], [49]. Estrogen itself has been shown to inhibit osteoclast formation and activity [50], [51]. Thus, the use of phenol red-free α-MEM was a logical choice for osteoclast studies.

The basal medium needed to be supplemented with a number of components that supported the formation of osteoclasts *in vitro* [4]. The complex process of fusion of monocytes during osteoclast differentiation made the process challenging. One of the components that appeared to be essential for osteoclast formation was 1α,25-dihydroxyvitamin D_3_ (vitamin D). It has been shown previously that culture of osteoclast progenitors in the presence of vitamin D led to an increased expression of integrin α_V_β_3_ on the cell surface [59]. This integrin is known to be expressed on the membrane of osteoclasts and their progenitors and to promote osteoclast progenitors’ attachment to the bone matrix through binding to vitronectin receptors and initiate osteoclast differentiation [25], [35], [59]. Besides, vitamin D was shown to enhance the formation of multinucleated TRAP-positive osteoclasts through increasing RANK expression in osteoclast progenitor cells. [34], [60]. RANK is the receptor to which RANKL binds, which is needed for osteoclast differentiation and activation [61]. In our study, adding vitamin D only slightly promoted the formation of multinucleated cells. This effect was probably through inducing the expression of the integrin α_V_β_3_ on monocyte surface and promoting the attachment and spreading of cells to the vitronectin coated surface. While most cells remained mononuclear, they seemed to migrate closer to each other, only missing a cue to fuse together. Vitamin D also increased TRAP activity even though most of the cells still seemed to be mononuclear monocytes. The TRAP release could be from mononuclear TRAP-positive cells which might be cells on the pathway to a multinucleated phenotype [62].

Cholesterol is one of the major components of biological membranes which affects their structure and function [63]. Cholesterol is insoluble in water, and it can be imported to the cells through LDL. LDL is a complex particle with a hydrophobic core of cholesterol and triglycerides which is surrounded by a hydrophilic membrane consisting of phospholipids, free cholesterol, and apolipoproteins [64]. LDL can enter cells via LDL receptors and upon its degradation, cholesterol can be released into the cytoplasm [64]. The regulation of cholesterol by LDL plays an important role in osteoclasts. It has been shown that depletion of LDL from serum impaired cellular membrane fusion events and osteoclast formation *in vitro*, while osteoclast formation was restored by adding LDL to the LDL-depleted serum [36], [37], [63]. In our study, when LDL was added as a supplement, multinucleated TRAP-positive cells were formed. Furthermore, TRAP activity of cells and their resorption capability indicated the formation of active osteoclasts in the complete serum substitute medium, with values comparable to FBS supplemented medium which contains not only LDL but also other types of lipoproteins such as high-density lipoprotein (HDL) [65], [66].

A major limitation of the current study is that FBS, an ill-defined cell culture supplement, has been used as the gold standard/control of the experiments. This means that the outcomes of the serum substitute medium have been compared to the outcomes collected from FBS containing medium which we stated ourselves could be unreliable due to the nature of the FBS and can be different in case of using different FBS batches/brands.

It should also be noted that the components of serum substitute medium did not induce osteoclastic differentiation in some of the investigated donors. The serum substitute medium might need further optimization. Various other molecules have been reported to have an influence on the differentiation of monocytes towards multinucleated osteoclasts which were not investigated in the current study, for example vascular endothelial growth factor (VEGF) [62], [67], [68], platelet-derived growth factor-BB (PDGF-BB) [69], [70], low concentrations (less than 10 nM) of dexamethasone [71], or epidermal growth factor (EGF) [72]. All these factors have been shown to promote osteoclast differentiation and survival. The benefit of adding any of these factors to our serum substitute medium should be investigated in future studies. Our new serum substitute medium for osteoclast cultures might help shed light on their precise functions on osteoclastogenesis of human monocytes without the overshadowing effect of the complex FBS composition.

## Conclusion

In this study, we have developed a serum substitute medium with potentials to promote the osteoclastogenesis of human PBMC derived monocytes. The developed serum substitute medium was able to support the formation of resorbing TRAP-positive multinucleated osteoclasts from human PBMCs derived monocytes, however it is still less optimal than medium containing FBS and needs to be more optimized in future studies. The serum substitute medium has the potential to eliminate the use of FBS when studying osteoclast differentiation and activity and provides an opportunity to study the effect of many other soluble factors on the bone resorption process without being dominated by unknown components of FBS.

## Acknowledgement

This work has been financially supported by the Dutch Ministry of Education, Culture and Science (Gravitation Program 024.003.013).

